# Transcriptome-based prediction of complex traits in maize

**DOI:** 10.1101/587121

**Authors:** Christina B. Azodi, Jeremy Pardo, Robert VanBuren, Gustavo de los Campos, Shin-Han Shiu

## Abstract

The ability to predict traits from genome-wide sequence information (Genomic Prediction, GP), has improved our understanding of the genetic basis of complex traits and transformed breeding practices. Transcriptome data may also be useful for GP. However, it remains unclear how well transcript levels can predict traits, particularly when traits are scored at different development stages. Using maize genetic markers and transcript levels from seedlings to predict mature plant traits, we found transcript and genetic marker models have similar performance. Surprisingly, genetic markers important for predictions were not close to or identified as regulatory variants for important transcripts. Thus, transcript levels are predictive not simply due to genetic variation. Furthermore, genetic marker models identified only one of 14 benchmark flowering time genes, while transcript models identified five. Our findings highlight that transcriptome data is useful for GP and can provide a link between traits and variation that cannot be readily captured at the sequence level.

## Introduction

The prediction of complex traits from genetic data is a grand challenge in biology and the outcome of such prediction has become increasingly useful for plant and animal breeding ^1,2^. Among the different approaches for connecting genotypes to phenotypes, Genomic Prediction (GP, or Genomic Selection) using all available markers was developed to overcome the limitations of Marker-Assisted Selection, which uses only significant quantitative trait loci (QTLs), for breeding traits that are controlled by many small effect alleles ^3,4^. Using GP, breeders are able to make data driven decision about what lines to include in their programs, speeding up and reducing the cost of developing the next generation of crops ^5,6^. Furthermore, because GP models are associating genetic signatures with phenotypes, untangling GP models has the potential to improve our understanding of the genetic basis of complex traits. However, as with related approaches such as genome wide association studies and QTL mapping, it remains difficult to go from associated genetic markers to the molecular basis for a trait ^7,8^.

There are a number of factors contributing to this difficulty. The variation in markers associated with phenotypes may not be the causal variants but are linked to the genes that control the trait in question. Considering that linkage disequilibrium distance can range from 1 kilobase (kb) in diverse maize populations ^9^ to ∼250 kb in *Arabidopsis thaliana* ^10^, the linked candidate genes can range from a few to a few hundreds. Even if the associated genetic variant is controlling the underlying phenotype, most variants associated with complex traits have small effect sizes and can be regulatory ^11^, which may not be linked to the genes they regulate. Furthermore, multiple regulatory variants that have indiscernible effects on their own, could interact epistatically to influence gene and ultimately trait expression. However, even with sufficient statistical power to detect genetic variants with small effect sizes and interactions between them, genetic information is connected to traits through multiple intermediate processes, including, for example, transcription, translation, epigenetic modification, and metabolism. Each of these intermediate processes represent an additional level of complexity that obscures the association between genetic information and a trait.

One solution is to account for these intermediate processes by integrating relevant omics data in addition to genetic variation. This approach has led to promising, but often mixed, results in plants. Current efforts have focused primarily on predicting hybrid performance using transcriptional information from the parental lines. For example, transcript level-based distance measures generated from transcripts associated with the trait were better than genetic markers in predicting hybrid performance in maize ^12,13^. However, when all transcripts were used (instead of a subset of pre-selected transcripts), model performance decreased ^14^. The performance of models based on transcript levels can be better or worse compared to those based on genetic markers depending on the trait. For example, transcriptome data performed better for predicting grain yield in hybrid maize populations, but genetic marker data performed better for predicting grain dry matter content in the same population ^15^. Similarly, in a maize diversity panel, GP models that combined transcript and marker data only outperformed models using markers alone for certain traits ^16^. Finally, efforts to integrate additional omic information to predict various traits in *Drosophila melanogaster* ^*17*^, and human diseases, such as breast cancer ^18^, and responses to treatment interventions, including acute kidney rejection and response to infliximab in ulcerative colitis ^19,20^, have demonstrated the potential usefulness of transcriptome data in the field of precision medicine.

Overall, these efforts provide reasonable evidence that transcriptome data could be useful for trait prediction. However, GP-based approaches that trained on the entire transcriptome data have not been used to better understand the genetic mechanisms for a trait. In addition, it is not known the degree to which transcriptomes obtained at a particular developmental stage can be informative for predicting phenotypes scored at a different stage. To address these questions, we used transcriptome data derived from maize whole seedling ^22^ to predict phenotypes (flowering time, height, and grain yield) at much later developmental stages. In addition to comparing prediction performance between genetic marker and transcriptome-based models, we also looked at whether transcripts and genetic marker features important for the prediction models were located in the same or adjacent regions. Finally, we determined how well our models were able to identify a benchmark set of flowering time genes to explore the potential of using GP to better understand the mechanistic basis of complex traits.

## Results and Discussion

### Relationships between transcript levels, kinship, and phenotypes among maize lines

Before using the transcriptome data for GP, we first assessed properties of the transcriptome data in three areas: (1) the quantity and distribution of transcript information across the genome, (2) the amount of variation in transcript levels, and (3) the similarity in the transcriptome profile between maize lines, with an emphasis on how these properties compared to those based on the genotype data. After filtering out 16,898 transcripts that did not map to the B73 reference genome or had zero or near zero variance across lines (see **Methods**), we had 31,238 transcripts. While the number of transcripts was <10% of the number of genetic markers used in this study (332,178), the distribution of transcripts along the genome was similar to the genetic marker distribution (**Fig. S1**). The log_2_-transformed median transcript level across lines ranged from 0 to 12.4 (median=2.2) and the variance ranged from 3×10^−30^ to 14.5 (median= 0.13), highlighting that a subset of transcripts had relatively high variation in transcript levels across maize lines at the seedling stage. To determine how similar transcript levels were between lines, we calculated the expression Correlation (eCor) between all pairs of lines using Pearson’s Correlation Coefficient (PCC). The eCor values ranged from 0.84 to 0.99 (mean=0.93). As expected, lines with similar transcriptome profiles were also genetically similar as there was a significant correlation between eCor values with values in the kinship matrix generated from the genetic marker data (Spearman’s Rank *ρ* = 0.27, *p* < 2.2×10^−16^; **Fig. 1A**). As a result, we were able to find clusters of lines that had both high transcript and genetic similarities (e.g. cluster a, b; **Fig. 1B, C**). However, most of the variation in eCor was not explained by kinship, which explained why we identified other clusters that had similar transcriptome profiles, but were not genetically similar (e.g. cluster c, **Fig. 1B, C**).

**Figure 1.**
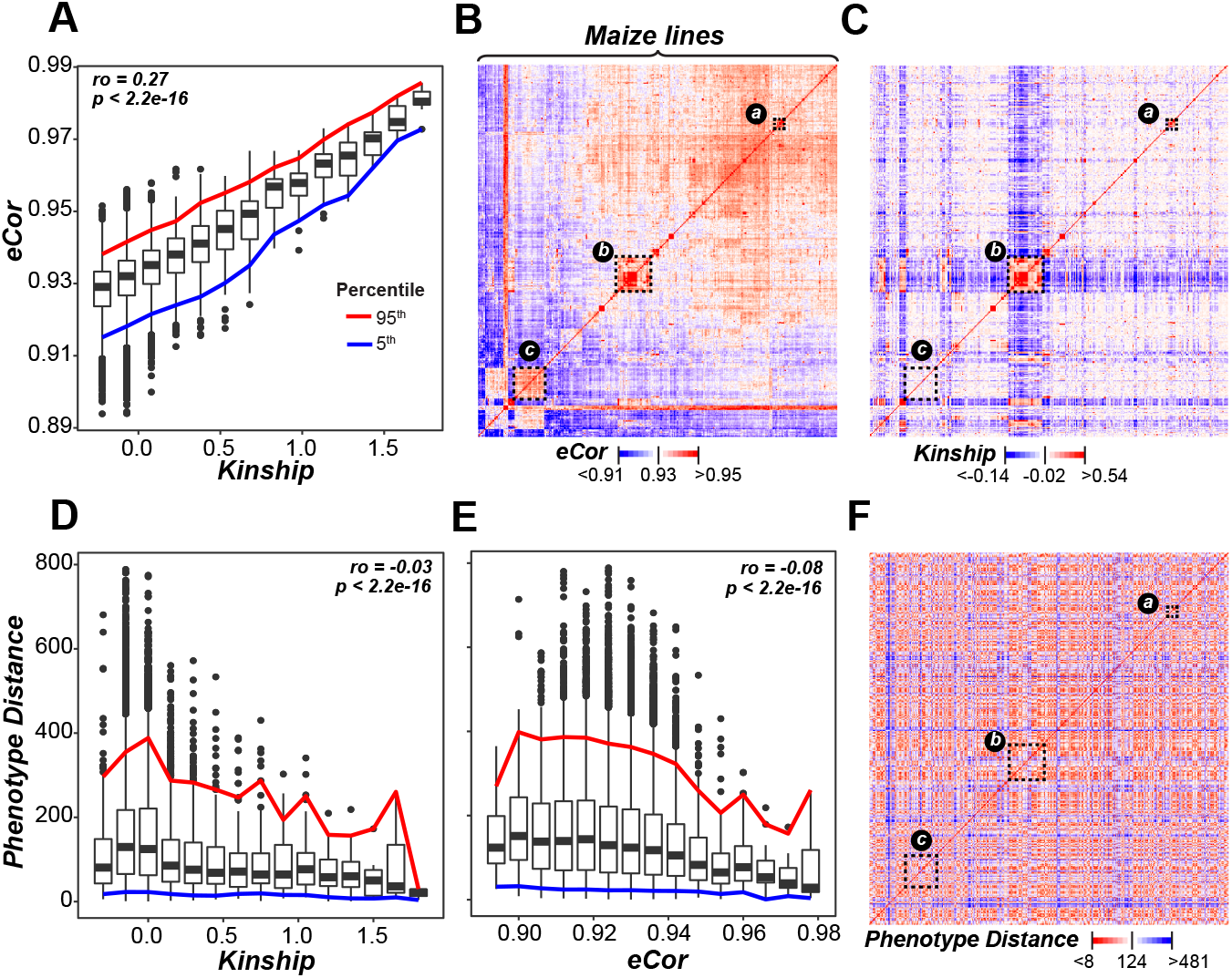
Relationship between lines from transcript and genetic marker data. **(A)** Relationship between kinship based on genetic marker data (X-axis) and expression correlation (eCor, in Pearson’s Correlation Coefficient (PCC)) based on transcript data (Y-axis). Boxplots show the median Y-axis value for each X-axis bin (sbin size=0.15) with the 5th (blue) and 95th (red) percentile range shown. The correlation between kinship and eCor was calculated using Spearman’s Rank Coefficient (ρ). **(B, C)** The relationships between lines based on eCor **(B)** or kinship **(C)** for all pairs of maize lines. Lines are sorted based on hierarchical clustering results using the eCor values. The blue, white, and red color scales indicate negative, no, or positive correlations, respectively. Dotted rectangles: indicating cluster of lines discussed in the main text. **(D, E)** The relationships between the Euclidian distance calculated with phenotype values (Phenotype Distance: Y-axis) and kinship **(D)**, and eCor **(E)**. Colored line: follow those in **(A)**. **(F)** The relationships between lines based on Phenotype Distance, where the lines were sorted as in **(B)**. Red: smaller distance (more similar). Blue: greater distances (less similar).

Because the basis of GP is to predict a phenotype from genetic data, we next asked if kinship or eCor were anti-correlated with the phenotypic distances between lines (see **Methods**). While both kinship (*ρ* = −0.03, *p* < 2.2×10^−16^; **Fig. 1D**) and eCor (*ρ* = −0.08, *p* < 2.2×10^−16^; **Fig. 1E**) were significantly, negatively correlated with the phenotype distance, the degree of correlation was minor. Furthermore, the groups of lines that clustered together based on their eCor (e.g. clusters a, b; **Fig. 1B, 1C**) did not have lower phenotypic distance (**Fig. 1F**). Taken together, these findings suggest that transcriptome data may be similarly informative as genotype data but capture difference aspect of phenotypic variation. We tested both of these interpretations further in subsequent sections.

### Predicting complex traits from transcript data

To test how useful transcriptome data was for GP compared to genetic marker data, we applied four approaches to predict three agronomically important traits in maize: flowering time, height, and grain yield. Because no one GP algorithm always performs best ^6,23^, we tested two linear algorithms (ridge regression Best Linear Unbiased Predictor (rrBLUP) and Bayesian-Least Absolute Shrinkage and Selection Operator (BL)), one non-linear algorithm (random forest: RF), and one ensemble approach (En; see **Methods**). To establish a baseline for our GP models, we determined the amount of the phenotypic signal that could be predicted using population structure alone, defined as the first five Principal Components from the genetic marker data. Then we built models for each trait using genetic marker data (G), kinship (K) derived from G, transcript levels (T), or expression correlation (eCor) derived from T (**Fig. 2**). Model performance was measured using PCC between the actual and the predicted phenotypic values. Across algorithms and traits, the K data resulted in models with the best predictive performance, while models built using the eCor data performed the worst (**Fig. 2, Table S1**). Furthermore, models built using G always outperformed models using T. Regardless, eCor and T-based models were significantly better than the baseline predictions (dotted blue line, **Fig. 2**), indicating transcriptome data can be informative in GP. Considering that the transcriptome is from seedling, it is particularly surprising that mature plant phenotypes can be predicted. Consistent with earlier findings ^24,25^, combining the predictions from multiple algorithms, known as an ensemble approach, resulted in the best predictive models (**Fig. 2**), and is therefore used to illustrate most of our findings.

**Figure 2.**
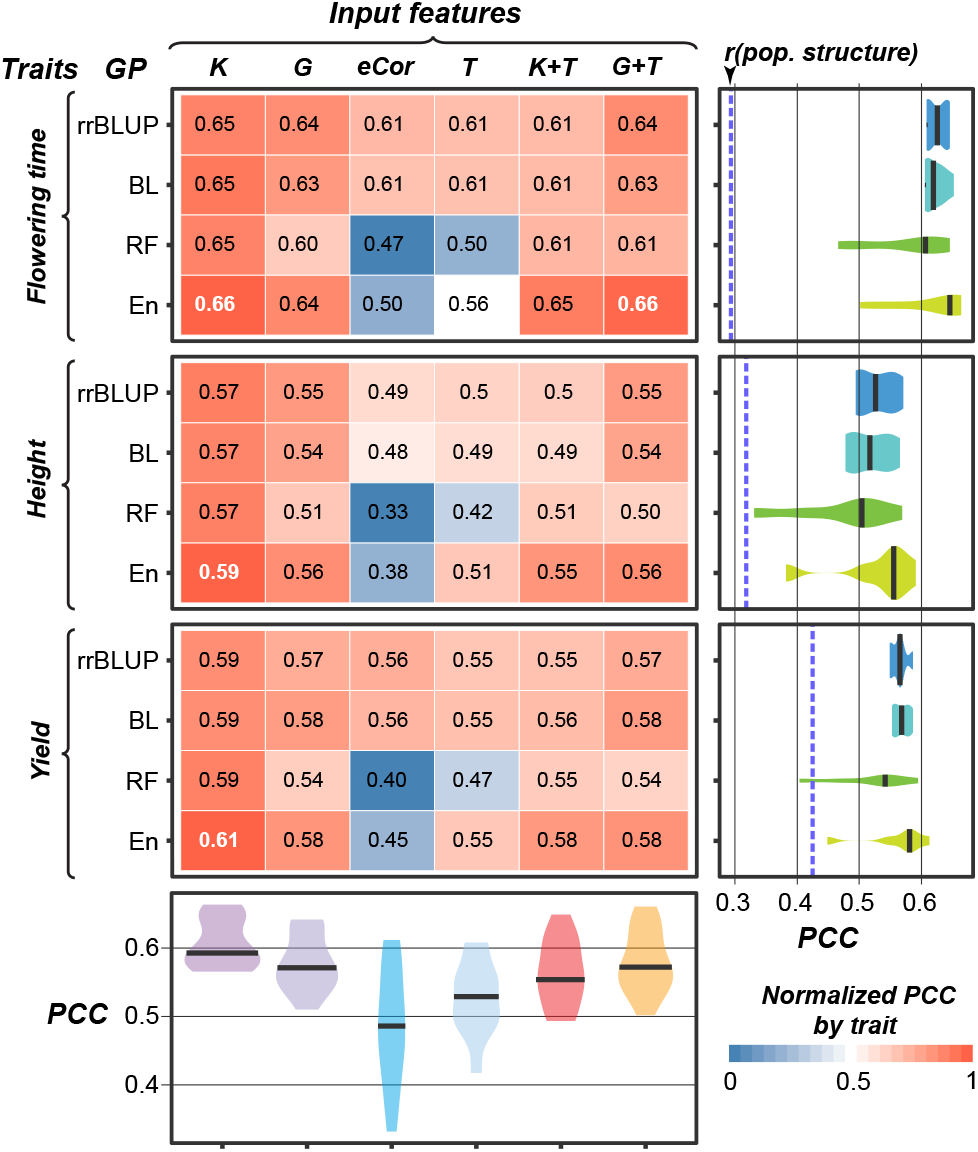
Genomic prediction model performance. PCCs between predicted and true values for three traits and four algorithms using six different input features. The darkest red indicate a normalized PCC of 1 (the algorithm/input feature combination performed the best for the trait), while the darkest blue has a normalized PCC of 0 (performed the worst. Original PCC values were shown in the boxes with the top performing model(s) in white. Right violin-plots show the PCC distributions among different input features for each algorithm (right). The median PCCs are indicated with black bars. The model performance PCCs based on only population structure are indicated with a blue dashed line. Bottom violin-plots show the PCC distributions among different algorithms for each input feature. rrB: ridge regression Best Linear Unbiased Predictor. BL: Bayesian Least Absolute Shrinkage and Selection Operator. RF: Random Forest. En: Ensemble.

Because the genetic marker and transcriptome data represented different types of molecular information that could be associated with the traits of interest, we hypothesized that their combination would be more informative and next built models that used combined data, either K+T or G+T. For most combined models, adding the transcript data did not significantly improve performance. The one exception was using RF to predict flowering time using G+T as input (**Fig. 2**). To assess if G or T data features tend to be more informative in predicting traits, we further quantified the importance score of each genetic marker and transcript feature for models using G+T data. The importance score represents the impact that each feature had on model performance defined according the algorithm used (see **Methods**). The importance scores assigned to transcripts in the G+T models were correlated with the scores from the T-only models (**Fig. S2A**), indicating that adding genetic marker features into the model did not impact the relative importance of transcript features. Because RF importance measures tend to be biased toward continuous features,^26^ we focused on rrBLUP and BL importance scores. For all three traits, the top 1,000 most important features were enriched for genetic markers relative to transcript features (Odds Ratio = 0.17 ∼ 0.44; all *p* < 1×10^−16^; **Fig. S2B**; **Table S2**). However, the top 20 most important features tended to be enriched for transcript relative to genetic marker features (Odds Ratio = 2.66 ∼ 13.0, *p* = 0.087 ∼ <1×10^−16^, **Table S3**), with transcript features making up the top two most important feature in all cases (**Fig. S2B**). The consistency with which transcript features were the most important for the models suggests that transcript information is useful for GP. Further highlighting its usefulness, when either the 200 most important transcripts or genetic markers were used to predict flowering time, models performed equally well (*r*=0.70 ± 0.010; *r*=0.71 ± 0.009, respectively).

### Comparison of the importance of transcripts verses genetic markers for model predictions

Because models built using transcript features outperformed baseline models based solely on population structure, we know transcriptome data contained information useful for explaining phenotypic variation. However, combining both datasets does not improve the model (K+T and G+T, **Fig. 2**), we hypothesized that this is because these two data types capture similar aspects of phenotypic variation. To address this, we assessed the extent to which the important genetic markers overlapped with or neighbored the genes where the important transcripts originated from (top; **Fig. 3A**). The genic region and flanking sequences within a defined window of an important transcript is referred to as the transcript regions (see **Methods**). For each trait and algorithm, we compared the importance assigned to the transcript with that of the genetic marker with the highest average importance in the transcript region (T:G pair).

**Figure 3.**
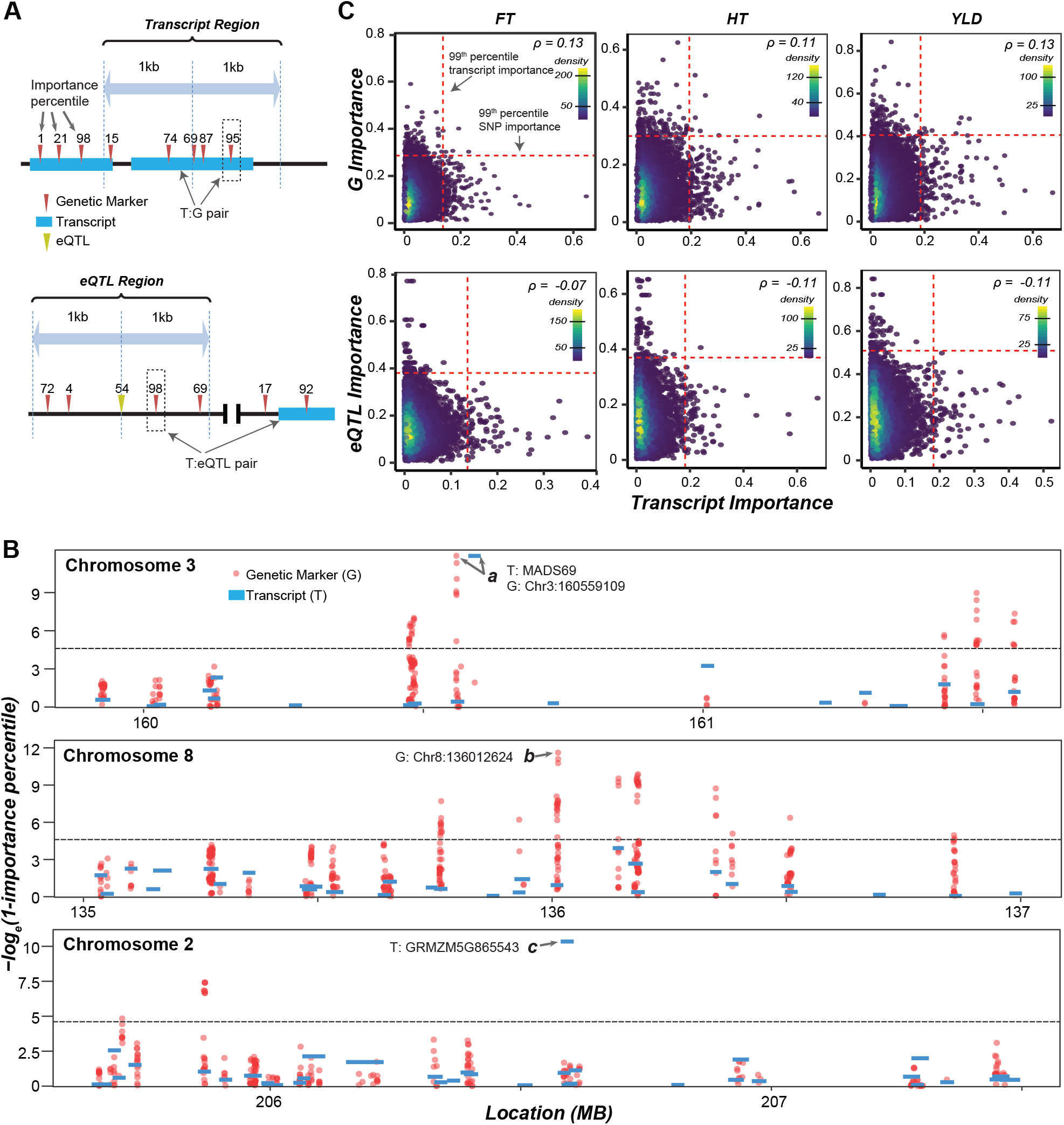
Correlation between genetic marker and transcript importance for flowering time. **(A)** Illustration of how transcript (T):genetic marker (G) (top graph) and T:expression Quantitative Trait Locus (eQTL) (bottom graph) pairs were determined. Genetic marker importance percentiles are shown above the genetic markers (red triangle) and eQTL (yellow triangle). A T:G pair was defined as the transcript and the most important genetic marker within the transcript region (top graph). A T:eQTL pair was defined as the transcript and the most important genetic marker within the eQTL region (bottom graph). **(B)** Manhattan plots of the transcript (blue bar) and genetic marker (red dot) importance scores (-loge(1-importance percentile)) in a 2Mb window surrounding top two genetic markers (top and middle plots) and transcripts (top and bottom plots) based on the Ensemble models for predicting flowering time. All genetic markers (i.e. not just the T:G pair) are shown. The threshold (gray dotted line) is set at the 99th percentile importance. **(C)** Density scatter plot of the importance scores (see Methods) of the genetic marker (Y-axis) and transcript (X-axis) for T:G pairs (top graphs) and of the eQTL genetic marker (Y-axis) and transcript (X-axis) for the T:eQTL pairs (bottom graphs) for three traits. The threshold (black dotted line) was set at the 99th percentile importance score for each trait and input feature type. The correlation between importance scores between transcript and genetic marker/eQTL pairs was calculated using Spearman’s rank (ρ).

Multiple window sizes were explored, and we used 2 kb (+/− 1kb from the center of a gene) where the feature importance correlation between transcripts and genetic markers was maximized (**Fig. S3**). Using this window size, 15,049 T:G pairs were identified. At the whole genome level there appeared to be regions where both genetic markers and transcripts were identified as important (**Fig. S4**). However, when we look closer, those regions mostly do not overlap. In some cases, the important genetic markers and transcripts were in linkage disequilibrium. Using the flowering time model as an example, we found the most important genetic marker was located within a gene upstream the most important transcript (GRMZM2G171650: *MADS69*; arrow a, **Fig. 3B**), but the two are in linkage disequilibrium ^*22*^. In most cases, there were no important genetic markers that were located nearby to important transcripts. For example, the second most important flowering time genetic marker was not located near important transcript regions (arrow b, **Fig. 3B**). Similarly, the second most important flowering time transcript (GRMZM5G865543) was over 0.6 Mb from an important genetic marker (arrow c, **Fig. 3B**). Across all traits and algorithms, T:G pairs were only moderately correlated (*ρ* = 0.09-0.13; **Fig. 3C, Fig. S5A**).

This lack of correlation is notable for the most important genetic markers and transcripts. For example, across the three traits, only 4-7 T:G pairs were both in the top 1% most important features from the ensemble models, and those pairs were never the top ranked genetic markers or transcripts from the model (**Fig. 3B)**. These findings argue against the notion that these two data types capture similar aspects of phenotypic variation as we hypothesized earlier. In light of this, we hypothesized that the lack of correlation was because important transcripts tend to be regulated by important *trans* factors located far beyond the transcript region. To test this, we assessed the degree to which important genetic markers identified as expression QTL (eQTLs) were associated with important transcripts. We identified 58,361 *cis* (62) and *trans* (58,299) eQTL associated with 7,052 transcripts and defined T:eQTL pairs for each of these transcripts by selecting the genetic marker within +/− 1kb of an eQTL for that transcript (i.e. eQTL region) with the highest average importance. Across all traits and algorithms, the importance of transcripts and eQTL in T:eQTL pairs was actually negatively correlated (*ρ* = −0.15 ∼ −0.06; **Fig. 3C, Fig. S5B**).

The lack of correlation between importance scores for T:G and T:eQTL pairs was in contrast to the relatively high correlation observed in feature importance between algorithms (*ρ*= 0.31-0.98), with rrBLUP and BL importance scores being the most correlated (*ρ*= 0.87-0.98) and the average correlation between genetic markers (*ρ* = 0.75) being higher than for transcripts (*ρ* = 0.55) (**Fig. S6**). Together with the findings that important genetic markers were not co-located and eQTL were not associated with genes that gave rise to the important transcripts for any of the three traits, these findings may suggest that transcriptome data is capturing layers of information, such as epigenetic signals, that are not captured by genome sequences. However, we cannot rule of the possibility that the eQTL approach is not sufficiently sensitive in identifying important *trans-*factors. Further study is needed to resolve these possibilities.

### Assessment of benchmark flowering time genes

Because the genetic basis for flowering time is well studied ^27–30^, we identified a set of 14 known flowering time genes (**Table S3**) and compared the ability of genetic marker and transcript-based models to reveal them as important using the T:G and T:eQTL pairs described earlier. Of the 14 benchmark genes, four had corresponding genetic markers in our T:G pair data. When we increased the flanking regions threshold to 20kb from the center of the transcript for defining T:G pairs, corresponding genetic markers were found for five additional benchmark genes. Two benchmark genes, *CCT1* and *PEBP4*, neither of which were members of a T:G pair, were associated with eQTLs. To account for differences in distribution and range of importance scores generated by different algorithms and numbers of features, the importance scores were converted to percentiles for comparison purposes.

Different benchmark genes were important (>95^th^ percentile) for models using the two different data types, with one and five benchmark gene considered important by the genetic marker-based and the transcript-based models, respectively (**Figure 4A**; **Table S4**). For example, the genetic marker located within the *RAP2* gene, which has been shown to be associated with flowering time in multiple studies ^22,31^, was identified as important based on genetic marker (99.7^th^-99.9^th^ percentile), but not transcript (59^th^-79^th^ percentile) data. In contrast, *MADS69, MADS1, PEBP24*, and *PEBP8* were identified as important using transcript data (95^th^-100^th^ percentile), but not using genetic marker data (16^th^-93^th^ percentile). Furthermore, with transcript data we were able to assess the importance of three genes (*ZAG6, REPB5, and PEBP2*) that were not located near genetic markers or associated with eQTL. For example, there were no eQTL associated with or genetic markers within the 40bp window of *ZAG6*, but *ZAG6* was identified as important (98^th^-99.9^th^ percentile) in the transcript-based models (**Fig. 4A**). For some of these benchmark genes, the region most closely linked to trait variation could be outside the +/− 20kb window. For example, as described above, the important genetic marker for *MADS69* (Chr3_160559109) is ∼32 kb upstream, but the two are in linkage disequilibrium ^22^ (see arrows in **a; Fig. 3B**). Taken together, these finding further highlight the usefulness of transcript data for identifying the genetic basis for variation in a trait.

**Figure 4.**
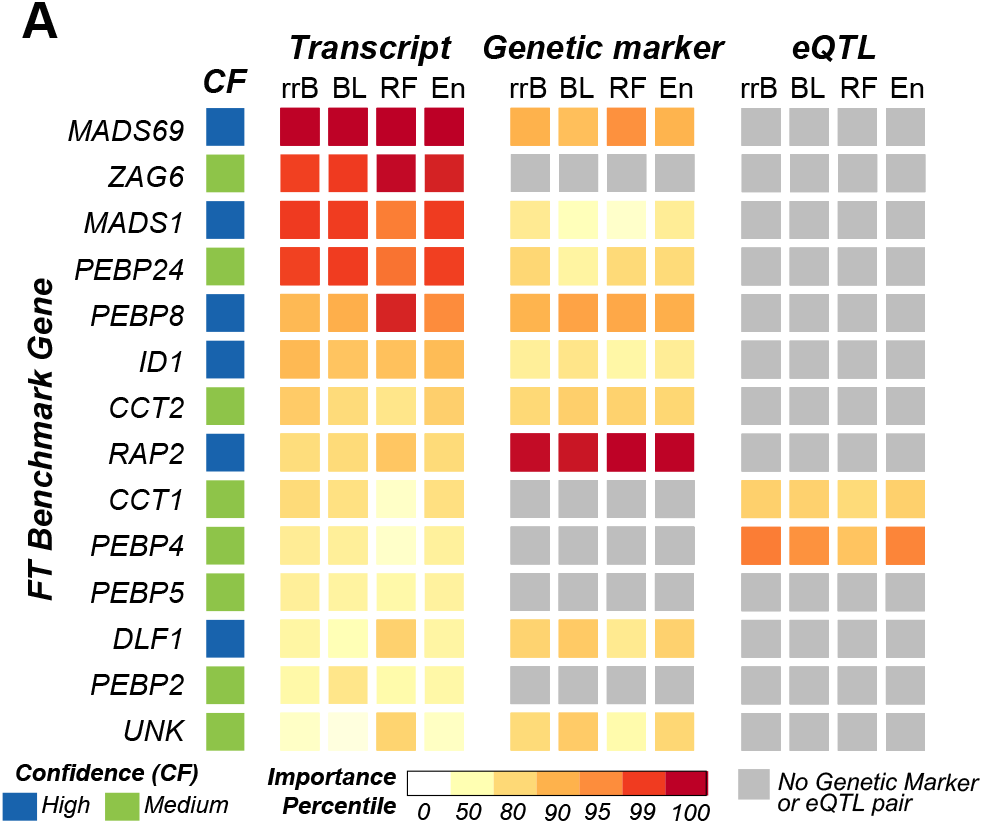
Comparison of transcript and genetic marker importance scores for benchmark flowering time genes. Importance percentile of each transcript (left) and genetic marker (right) pair as determined by each of the 4 algorithms (X-axis). Genes are sorted based on hierarchical clustering of their importance percentiles. Gray boxes designate benchmark genes that did not have genetic markers within a 40kb window. Confidence levels (high or medium) were assigned based on the type of evidence available for the benchmark gene (see Methods). Algorithms were abbreviated as in Figure 2.

### Improving our understanding of the genetic basis of flowering time using transcriptome data

An open question was why transcript-based models were able to identify five benchmark flowering time genes as important that were not identified by genetic marker-based models and if transcriptome data could be used to better understand the genetic basis of flowering time. To understand why benchmark genes were not uniformly identified as important for flowering time when using both genetic marker and transcript data, we determined the extent to which transcript levels and the genetic marker allele (i.e. major or minor) were related to flowering time. As expected, we observed the most significant differences in flowering time for the transcripts (**Fig. 5A, Fig. S7A**) and genetic markers (**Fig. 5B, Fig. S7B**) that were identified as important by our models. For example, *MADS1* was important only in the transcript-based models and transcript level was significantly correlated with flowering time (*p* = 0.0001; **Fig. 5A**). In contrast, lines with the major allele for the genetic marker that paired with the *MADS1* transcript (Chr9: 156980141) did not flower at a significantly different time than lines with the minor allele (*p* = 0.062; **Fig. 5B**). Another example was *RAP2*, which was important only in the genetic marker-based models. Lines with the major allele in *RAP2* were more likely to flower late (*p* < 1×10^−4^), but *RAP2* transcript levels did not significantly correlate with changes in flowering time (*p* = 0.33). Overall, benchmark genes were more likely to have transcript levels associated with flowering time (**Fig. 5C**) than genetic marker alleles associated with flowering time (**Fig. 5D**).

**Figure 5.**
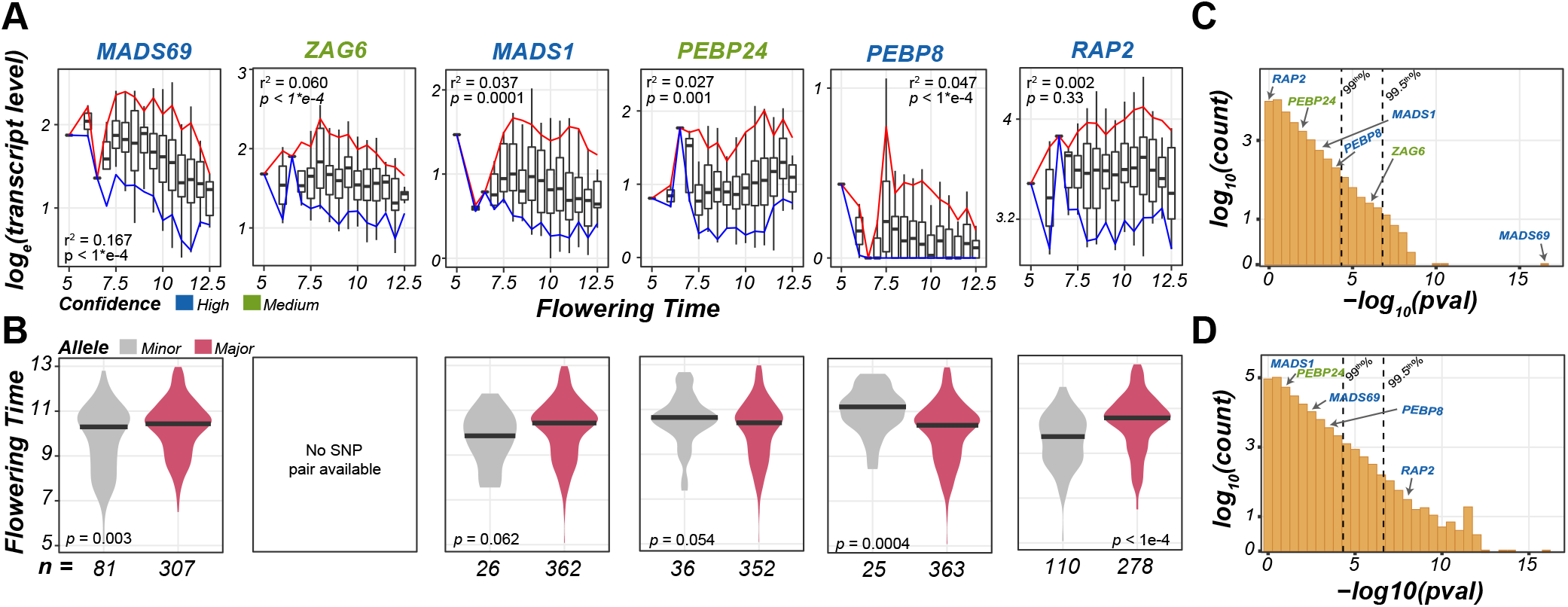
Relationship between transcript level/allele type and flowering time for benchmark genes. **(A)** Boxplots show the transcript levels (loge(Fold-Change)) over flowering time bin with the 5th (blue) and 95th (red) percentile range shown. Flowering time was defined as the growing degree days/100. Linear models were fit and adjusted r2 and p-values are shown. Confidence levels of benchmark genes were designated as in **(4)**. **(B)** Distributions of flowering time for lines with the major (red) or minor (gray) alleles for the genetic marker paired with each benchmark gene as indicated in **(A)**. Differences in flowering time by allele were tested using t-tests. **(C)** Number of transcripts (Y-axis) for which transcript levels were associated with flowering time in linear models within p-value bins (-log10(p-value); X-axis). Benchmark genes are labeled as in **(A)**. **(D)** Number of genetic markers (Y-axis) for which differences in flowering time by allele from t-tests were within p-value bins (-log10(p-value); X-axis). Benchmark genes are labeled as in **(A)**.

Importantly, using the transcriptome data we were also able to understand in more detail the impact of the benchmark genes on flowering time. For example, variation in transcript levels of *MADS69* accounted for 16.7% of the variation in flowering time, more than any other transcript, where lines with lower levels of transcription flowered later. Modulation of *MADS69* expression levels has recently been patented as an approach to controlling flowering time^32^. Similarly, *MADS1* transcript levels explained 3.7% of the variation in flowering time, with lines with lower levels of transcription flowering later. This is consistent with what has been observed experimentally, where down-regulation of *MADS1* results in delayed flowering time ^33^. For medium confidence benchmark genes (i.e. identified through association studies), the specific roles of the genes on flowering time are not well understood, but by finding positive or negative correlations between transcript levels and the underlying phenotypes, more mechanistic details can be interred. For example, transcript levels of *ZAG6* had the second largest impact on flowering time, accounting for 6% of variation, with increased transcript levels associated with earlier flowering. Another example is *PEBP24*, with transcript levels of *PEBP24* accounting for 2.7% of the variation in flowering time. Unlike many of the other benchmark genes, increased *PEBP24* transcript levels were associated with later flowering time. Overall, the identification of these medium confidence benchmark genes as important transcript indicates the relevance of transcriptional regulation in their flowering time functions.

While using the benchmark genes allowed us to assess the usefulness of transcript levels compared to genetic marker information for identifying genes involved in flowering time, we should note that many non-benchmark genes were also identified by our models as important. For example, from the Ensemble model, there were 154 important, non-benchmark transcripts with importance scores falling between the two most important benchmark genes (*MADS69*, 100^th^ percentile; *ZAG6*, 99.5^th^ percentile; yellow, **Table S5**). While seven of those in between transcripts were annotated with the Gene Ontology (GO) term “flower development” (GO:0009908, green, **Table S5**), these 154 non-benchmark transcripts were not enriched for this GO term (*q* = 1.0). In fact, neither these transcripts nor any other set of important transcripts from models based on other algorithms (see **Methods**) were enriched for any GO terms. Therefore, from our transcript-based GP models we have identified 147 high ranking transcripts, many of which have unknown functions, that are among the most important in predicting flowering time in maize but do not play known roles in this process. For example, both GRMZM5G865543 and GRMZM2G023520, the second and third most important transcripts respectively from the Ensemble model, are unknown genes. Note that the transcriptome data is from the seedling stage. It is possible that genes of these important transcripts influence biological processes in earlier stage of development that influence flowering time later. To further our understanding of the genetic basis of flowering time control and the connections between juvenile and adult phenotypes, these important transcripts are prime candidates for future genetic studies.

## Conclusions

We have generated predictive models that use genetic markers, transcripts, and their combination to predict flowering time, height, and yield in a diverse maize population. While models built using transcriptome data did not outperform models that used genotype data, transcript-based models performed well above random expectation, and in many cases, performance was similar to that of genotype-based models. We found that transcripts and genetic markers from different genomic regions were identified as important for model predictions. Furthermore, by assessing the relative importance of the features used to build the models, we found that transcript-based models identified more known flowering time associated genes than genetic marker-based models. These findings underscore the usefulness of transcript data for improving our understanding of the genetic mechanisms responsible for complex traits.

There are four possible mechanistic explanations of why transcript levels could have a similar predictive power as genetic markers. First, *cis-*regulatory variants that impact transcript levels, are all more likely to be similar between closely related individuals. Therefore, the ability of transcript data to predict phenotypes is simply a reflection of that dependency. However, we demonstrated that the most informative transcript features for predicting maize phenotypes are distinct from the most informative genetic marker features found in the transcript regions. While for some important transcripts, the associated important genetic marker could be in linkage disequilibrium but outside of the 2kb window used in our study (e.g. ∼32 kb away in the case of *MADS69*), overall as we increased the transcript region window size, the correlation between the importance scores assigned to T:G pairs decreased, suggesting this is not generally the case. Thus, the second explanation is that there are *trans-*regulatory variants, e.g. due to transposon polymorphisms or transcriptional regulators, that play a major role. However, we found that the importance of eQTLs (99.9% *trans*) and their associated transcripts were not positively correlated, suggesting that the *trans*-regulatory variation we identified cannot explain why transcript variation is predictive of phenotypic variation either. However, considering the challenges in identifying eQTLs due to mixed tissues used ^34^ and in modeling epistatic interactions ^35^, we cannot conclusively rule of this possibility. The third explanation is that transcription is a molecular phenotype caused by the integration of multiple genetic marker signals, both *cis* and *trans*, that may not have had strong signals individually. The fourth explanation is that there are epigenetic variants contributing to expression variation. It remains to be determined what the contribution of epigenetic variation is on our ability to use transcript data to predict phenotypes.

One surprise is that the transcript data generated during the V1 seedling stage on whole seedlings can predict adult plant phenotypes. We reason that complex traits, such as flowering time, are influenced by more than just canonical genes that act immediately prior to the growth and developmental sequences leading to flowering. For example, early developmental events such as cotyledon damage ^36^, root restriction ^37^, and photoperiod and temperature changes ^38^ can impact flowering time in mature plants. Therefore, early development transcript differences could eventually result in different flowering time. However, we anticipate that if transcript data collection occurred temporally and/or spatially closer to the phenotype data the predictive power of transcript levels would increase, and likely perform better than genetic marker-based models. Finally, an area of active research in GP is the incorporation of Genotype by Environment (GxE) interactions into predictive models ^39–41^. One potential benefit of using transcript information for GP could be that GxE interactions would be picked up by transcript level signals. Because transcriptome data used in our study was from whole seedlings (i.e. not the same individuals that were phenotype), this could not be tested.

Our findings highlight an important benefit of using transcript data to better understand the genetic basis of a trait. While it can be difficult to associate signals from a number of small effect genetic markers or even a single large effect genetic marker back to a specific gene, transcript level information is inherently associated with genes. Because of the importance of regulatory variation on complex traits ^11^, the use of transcript information in GP could be crucial for deciphering the contribution of regulatory variation to the genetic basis of traits. Therefore, while we observed that in terms of predictive ability, genetic marker data outperformed transcript data, expression differences are more straightforward to interpret than sequence polymorphisms. In practice, this meant that transcript-based models identified five benchmark flowering time genes, while genetic marker-based models only identified one and it highlighted our finding that more insight into the genetic basis of complex traits can be gained when transcriptome data are considered.

## Author Contributions

C.B.A., J.P., and S.-H.S. conceived and designed the study. J.P. assembled the data. C.B.A. wrote modeling code and the main manuscript. C.B.A. and J.P. ran the models. All authors assisted with interpretation of results and manuscript editing.

## Acknowledgements

We thank Richard Amasino, Wolfgang Busch, and David Lowry for their help in interpreting our findings. This work was partly supported by NSF Graduate Research Fellowship (Fellow ID: 2015196719), Graduate Research Opportunities Abroad (GROW) Fellowship to C.B.A.; the U.S. Department of Energy Great Lakes Bioenergy Research Center (BER DE-SC0018409) and National Science foundation (IOS-1546617, DEB-1655386) to S.-H.S.

## Methods

### Genotypic, transcriptomic, and phenotypic data processing

The phenotypic ^42^, and genotypic and transcriptomic ^22^ data used in this study were generated from the pan-genome population consisting of diverse inbred maize lines. Genotype, transcriptome, flowering time, height, and yield data was all available for 388 lines out of the 503 maize pan-genome panel and were used for the study (**Table S6**). Genetic marker scores derived from RNA-seq reads were converted to a [-1,0,1] format corresponding to [aa, Aa, AA] with the more common allele (AA) designated as 1. The genetic marker positions were converted from maize B73 reference genome A Golden Path v2 (AGPv2) to AGPv4.37. The AGPv2 genetic markers that did not map to AGPv4.37 and genetic markers with a minor allele frequency less than 5% were removed, resulting in 332,178 genetic markers.

Transcriptomic data from whole-seedling tissue including root at the V1 stage from ^22^ was processed to remove loci that did not map to AGPv4.37. The remaining maize B73 genes were filtered with default settings of the nearZeroVar function from the R caret package to remove genes with zero or near zero variance (> 95% of the lines sharing the same transcript level) across lines. After the filtering steps, transcript counts for 31,238 genes were retained in the final dataset. The raw transcripts per million count data were transformed with a log_e_ + 1 transformation before the data were used in subsequent analyses. To assess if transcriptome data had predictive power beyond random expectation, transcriptome data was permuted by gene, so each gene had the same distribution of transcript values, but the values were randomly assigned to different maize lines for building the transcriptome shuffled models. To compared important transcripts and genetic markers from GP models, transcripts were converted from AGPv3 to v4, only genes with one to one correspondence between AGPv3 and v4 were included in this analysis.

### Comparison of transcript and genetic marker data

Three different approaches were used to determine the similarity between lines based on the three different data types. For the genotype data, a kinship matrix was generated using the centered Identity By State (IBS) method ^43^ implemented in TASSEL v5.20180517 ^44^. For the transcript data, we generated an expression Correlation (eCor) matrix by calculating the Pearson Correlation Coefficients (PCCs) of transcript values between lines using the cor.test function in R. The eCor matrix was normalized between 0 and 1 and the diagonal was set as 1. Finally, for phenotype data, we calculated the Euclidean distance between lines using the distances package in the R environment. The correlation between kinship, eCor, and Phenotype Distance between pairs of lines was calculated using PCC.

### Genomic prediction models and model performance

Because part of the phenotypic signal observed in GP models may be due to population structure within the breeding population, we established a baseline for our GP models by using the first 5 principle components generated using the marker data alone, to predict phenotype values. Four methods were used for each trait, two linear-parametric methods: ridge regression-Best Linear Unbiased Predictor (rrBLUP)^45^ and Bayesian Least absolute shrinkage and selection operator (BL)^46^, and one non-linear and non-parametric method: Random Forest (RF)^47^, and one ensemble based approach (En)^48^. Both rrBLUP and BL were implemented in R using the “rrBLUP” and “BGLR” packages respectively. RF was implemented in python using Scikit-Learn ^49^. Ensemble predictions were generated by taking the mean of the predicted trait values from rrBLUP, BL, and RF. A grid-search was performed on the first 10 of the 100 cross-validation replicates to find the best combination of parameters for the RF model. Parameters tested included max tree depth (3, 5, 10, and 50) and the max number of features included in each tree (10%, 50%, 100%, square root, and log_2_).

The predictive performance of the models was compared using the PCC. The PCC between the predicted (Ŷ) and the true trait value (Y) and was computed using the cor() function in R for rrBLUP and BL or the NumPy corrcoef function in Python for RF. One hundred replicates of a five-fold cross validation approach were applied to maximize the data available for model training without resulting in overfitting. For each replicate, the lines were randomly divided into 5 subsets, where each subset is used as the testing set once and the rest 4 subsets combined to train the model, resulting in a total of 500 cross-validated runs. PCC was calculated using only the predicted values from the testing set for each run.

### Genetic marker/transcript importance analysis

In order to identify features important for building the predictive models, feature importance information was extracted from each model established with one of four methods: rrBLUP, BL, RF, and Ensemble. For rrBLUP, the importance metric was the marker effect ($u) calculated by mixed.solve in the R rrBLUP package. For BL, the importance metric was the estimated posterior mean ($ETA) calculated using the R BGLR package. The absolute value of marker effect and estimated posterior mean were used since the features are categorical with no particular meaning for the sign of importance metrics. For RF, the importance metric was the Gini importance, collected using the _importance_score function built into the Scikit-Learn implementation of RF. The Gini importance is the total decrease in node impurity (i.e. the homogeneity of classes in a node) after a particular feature is used to split a node. Node impurity decreases as instances from one of the classes are removed from the node, leaving a greater proportion of instances from the other class. Importance metrics from rrBLUP, BL, and RF were averaged over the 100 cross-validation replicates. Ensemble importance scores were calculated by normalizing the average importance scores from each model and each method between 0 and 1, then taking the mean of normalized importance scores across the three algorithms. Enrichment for transcript compared to genetic marker features within the top 1000 or top 20 features was done using Fisher’s Exact Test, where the number of transcript features in and not in the top X features was compared to the number of genetic marker features in and not in the top X features.

To determine the degree to which the importance of a transcript correlates with the importance of nearby genetic markers, the genetic marker *G* with the greatest mean importance score within a fixed window from the center of a genomic region *R* where a transcript *T* mapped to was selected among genetic markers in region *R*, referred to as a T:G pair (**Fig. 3A**). To identify the effect of window size, a series of window sizes ranging from 1-40kb were tested. For each window size, the Spearman’s Correlation (*ρ*) was calculated between the importance scores of T:G pairs. The window size with the highest correlation (2kb) was chosen (**Figure S4**). For this analysis, transcripts without location information or without one-to-one mapping between AGP V3 to V4 were removed, leaving 24,412 transcripts. With a window size of 2kb, additional transcripts were dropped because there was not a genetic marker within that window, resulting in 15,049 transcripts to be included in the downstream analysis.

To determine the degree to which the importance of a transcript correlated with the importance of *trans-*regulatory variants, significant eQTLs (multiple testing corrected *p* < 0.05) were identified for each transcript using the linear regression (modelLINEAR) approach from MatrixeQTL implemented in R. Benjamini-Hochberg false discovery rate correction was used to adjust *p* for multiple testing and eQTLs were considered significant if adjusted *p* < 0.05. The distance for considering eQTL as *cis* was 1 mega base ^50^, however, because <0.1% of eQTL identified were *cis*, all eQTL were analyzed together. The importance of an eQTL or the neighboring genetic marker located within a 2kb window of the eQTL with the greatest average importance score was compared to the importance of the transcript with the eQTL in question (T:eQTL pair).

Enrichment of Gene Ontology (GO) terms associated with important transcripts compared to the reference genome was tested using agriGO v2 ^51^. The enrichment *p*-values are corrected for multiple testing by agriGOv2 using FDR. The top 10, 25, and 100 transcripts from each algorithm, excluding the benchmark flowering time genes, were tested against the reference genome. Additionally, the top 153 transcripts, excluding benchmark genes, from the ensemble algorithm and the union of the top 10, 25, and 100 transcripts from all four algorithms were tested.

### Benchmark flowering time genes

We compiled a list of genes known to be involved in flowering time based on evidence from knockdown experiments ^27–30,33^ and/or association study ^22,52^. Genes were assigned confidence levels based on the type of evidence available, with experimental evidence considered high confidence, association study evidence and significant similarity with known flowering time genes from other species considered medium confidence (**Table S8**). Because some of these genes did not have genetic markers located within the 2kb window of the center of the transcript, progressively larger windows were used to identify the most important nearby genetic marker up to 40kb. To compared importance scores across algorithms and between models using G or T data as input, percentiles were used. To determine if transcripts or genetic markers assigned to flowering time benchmark genes were associated with flowering time in this study, linear models and t-tests, respectively, implemented in R were used.

### Data Availability

All data and code needed to reproduce the results from this study is available on GitHub including genomic, transcriptomic, and phenotype data (https://github.com/ShiuLab/Manuscript_Code/tree/master/2019_expression_GP/data), codes to run rrBLUP and BL models (https://github.com/ShiuLab/GenomicSelection), codes to run RF models (https://github.com/ShiuLab/ML-Pipeline), as well as R code used for preprocessing, T:G/eQTL pairing, eQTL analysis, and additional statistical analyses (https://github.com/ShiuLab/Manuscript_Code/tree/master/2019_expression_GP/scripts).

## Supplemental Figures

***Figure S1. Distribution of genetic marker and transcript data across maize chromosomes***

Number of genetic markers (top) and transcripts (bottom) included in this study in 1 Mb bins across the maize chromosomes.

***Figure S2. Feature importance analysis for G+T models***

**(A)** Relationships between importance scores for transcripts from the T (X-axis) and G+T (Y-axis) flowering time prediction models established with rrBLUP (left column), BL (middle column), and RF (right column). The Pearson’s Correlation Coefficient (r) is shown in the top left corner. **(B)** Distribution of importance scores for the top 1,000 (inset = top 20) features from the G+T models for three traits using rrBLUP (top row) and BL (bottom row). Transcript features are in purple and genetic marker features are in yellow.

***Figure S3. Impact of transcript region sizes on importance correlation between transcript:genetic marker pairs***

The correlation (green) between importance scores for transcript:genetic marker pairs and the number of pairs found (blue) as the transcript region size increases. Shown here are the correlation scores when using top (solid) or the 95th percentile (dashed) mean importance score of genetic markers in the transcript region.

***Figure S4. Manhattan plot of importance scores from Genomic Prediction models***

Manhattan plots of genetic marker (top) and transcript (bottom) importance scores for predicting **(A)** flowering time, **(B)** height, and **(C)** yield. Threshold importance scores (dotted blue) were set at the 99^th^ percentile importance score for each trait, algorithm, and input feature type (i.e. genetic markers or transcripts). Genetic markers and transcripts falling above that threshold colored in blue.

***Figure S5. Correlation between genetic marker/eQTL and transcript importance***

Density plot of the importance scores of **(A)** genetic markers (G, Y-axis) and transcripts (T, X-axis) from T:G pairs and **(B)** eQTL (eQTL, Y-axis) and transcripts (T, X-axis) from T:eQTL pairs. The threshold was set (red dotted line) as the 99^th^ percentile of the normalized importance score for each trait, algorithm, and input feature type. The correlation between transcript and genetic marker importance was calculated using Spearman’s Rank (*ρ*).

***Figure S6. Correlation between feature importance between algorithms***

Density scatter plot of the importance scores of genetic markers (top) and transcripts (bottom) generated with rrBLUP and BL (left), rrBLUP and RF (middle), as well as BL and RF (right). The correlation between importance scores between algorithms was calculated using Spearman’s Rank (*ρ*).

***Figure S7. Relationship between transcript levels and alleles and flowering time for benchmark genes***

(A) Boxplots show the median transcript level (log(Fold-Change)) for each flowering time (Growing Degree Days (GDD)/100) bin with the 95th (red) and 5th (blue) percentiles shown. Linear models were fit and adjusted r^2^ and p-values are shown. (B) Violin-plots of the distribution of flowering time (GDD/100) for lines with the major (blue) or minor (gray) allele for the genetic marker paired with each benchmark gene. Significant differences in the GDD by allele were tested for using t-tests.

## Supplemental Tables

***Table S1. Model performance by feature input type and algorithm***

***Table S2. Enrichment of transcript vs. genotype features among the top most important features from G+T models***

***Table S3. Description of benchmark flowering time genes, including evidence for flowering time association and T:Gs and T:eQTL pair information***

***Table S4. Importance scores and percentiles for benchmark gene transcripts, and genetic marker and eQTL pairs***

***Table S5. Top 1000 most important transcripts for flowering time from the Ensemble models***.

***Table S6. Account of data (Genetic Marker, Transcript, Phenotype) availability for maize lines and decision to include line in the study***

